# Targeting GPR183 to reduce peripheral sensitization: evidence from rodent and human tissue analyses

**DOI:** 10.64898/2026.01.15.699745

**Authors:** Kyle Oberkrom, Kathryn Braden, Evelyn Li, Samay R. Shah, Dragan Maric, Andrew J Mannes, Paola Pacifico, Nirupa D Jayaraj, Christopher K. Arnatt, Daniela Menichella, Matthew R. Sapio, Daniela Salvemini

**Author notes:** Corresponding author: Daniela Salvemini;. Saint Louis University School of Medicine, Pharmacology and Physiology, 1402 S. Grand Blvd., St. Louis, Mo. 63104. **Authorship Contributions**: *Participated in research design:* Oberkrom (designed mechanical allodynia, nitric oxide synthase, cyclooxygenase, knockout, and rat incision experiments), Braden (designed GPR183 activation in thermal sensitivity and inhibition with U0126 for mechano-and thermal hypersensitivities), Sapio (designed in situ hybridization and human experiments, contributed to research framing), Salvemini (directed the project). Conducted experiments:* Oberkrom (performed mechanical allodynia, nitric oxide synthase, cyclooxygenase, knockout, and rat incision experiments), Braden (performed Hargreave’s methods and von Frey for U0126 effects on GPR183-induced mechano-allodynia), Li (Performed in situ hybridization and tissue preparation), Maric (performed multicolor microscopy, image stitching, and composition). Pacifico and Jayaraj (performed DT injections, mechanical allodynia test and histology of LCs). Provided reagents:* Arnatt (synthesis of compounds). Performed data analysis:* Oberkrom (analyzed GPR183-induced mechanical allodynia, nitric oxide synthase, cyclooxygenase, knockout, and rat incision behavioral and protein expression data), Braden (analyzed thermal sensitivity – Hargreaves – data and mechanical sensitivity data following U0126 treatment), Sapio + Shah (Analyzed RNA-Seq datasets and assessed in situ hybridization data). Pacifico, Jayaraj and Menichella (analyzed LC-ablation mechanical allodynia data and quantified LC density). Wrote or contributed to writing of the manuscript:* Oberkrom, Shah, Pacifico, Menichella, Sapio, Salvemini. Authors claim no conflict of interest.

## Abstract

Peripheral sensitization is a key process in the development of painful inflammatory conditions, driven in part by immune-cell mediator release following tissue injury. The G protein–coupled receptor, GPR183, predominantly expressed on immune cells, regulates their migration, positioning, and mediator production. Yet its role in peripheral sensitization and the specific immune cells involved remains insufficiently understood.

In rats, intraplantar injection of 7α,25-dihydroxycholesterol (7α,25-OHC), the most potent endogenous GPR183 ligand, produced long-lasting nociception that was prevented by the selective GPR183 antagonist SAE-14. Because GPR183 activates ERK signaling, which influences pain pathways including nitric oxide synthase (NOS) activity and NO formation, we used NOS inhibitors and knockout animals to test the contribution of inducible and neuronal NOS isoforms to 7α,25-OHC-induced sensitization. We found that both isoforms influence this response, independent of cyclooxygenases. In a well-characterized rat incisional injury model, GPR183 protein expression increased in injured paw tissue, and SAE-14 reversed hypersensitivity. Meta-analysis of human post-surgical skin samples similarly showed elevated GPR183 expression and transcriptional changes favoring 7α,25-OHC production after injury. We identified macrophages and Langerhans cells (LCs) as the principal GPR183-expressing cell types in human skin. LC ablation studies revealed that 7α,25-OHC-evoked hypersensitivity does not depend on LCs, implicating GPR183^+^ macrophages as predominant drivers of GPR3-induced hypersensitivity. Overall, our findings define the cellular and molecular pathways linking GPR183 to peripheral sensitization and highlight GPR183 antagonism as a promising strategy for pain management.

## 1. Introduction

Peripheral sensitization is characterized by a reduced threshold or heightened responsiveness of sensory afferents to noxious stimuli (i.e., hyperalgesia) or the perception of non-noxious stimuli as painful (i.e., allodynia), that commonly arise following tissue injuries, such as trauma and inflammation. Immune cells contribute to this process by releasing cytokines, chemokines, and lipid mediators to isolate tissue damage and support recovery [2; 85]. Hypersensitivities typically resolve alongside inflammation; however, extended hypersensitization can evolve into significantly burdensome chronic pain states [40].

Oxysterols are endogenous, bioactive cholesterol derivatives implicated in inflammation [26; 32; 63]. For example, 7α,25-dihydroxycholesterol (7α,25-OHC) exacerbates lung inflammation via the G_αi_-coupled receptor GPR183 [26; 70]. GPR183 is a chemotactic receptor expressed on most immune cell lineages, including B cells [7], macrophages [26; 57], dendritic cells [18; 30; 79], eosinophils [70], T cells [27], and Langerhans cells (LCs) [16; 60]; yet, various datasets have shown that GPR183 is most highly expressed by microglia, macrophages, and LCs [38; 51; 60; 75]. In these cells, GPR183 modulates migration and positioning along a gradient of its most potent ligand, 7*α*,25-OHC [3; 32; 70]. Additionally, GPR183 promotes cytokine and chemokine expression changes that favor nociception by reducing anti-inflammatory and analgesic pathways while inducing inflammatory and noxious signaling [3; 9; 18; 26]. We implicated GPR183 in central sensitization where it signals through mitogen-activated protein kinase (MAPK)/extracellular signal-regulated kinase (ERK), developed highly selective GPR183 antagonists, and highlighted their potential as novel, non-opioid based analgesics [9; 10].

While GPR183 has been explored in central sensitization, its role in peripheral sensitization is limited to one investigation [58].

Nitric oxide synthase (NOS) generates nitric oxide (NO) from L-arginine [35], which is rapidly synthesized following tissue injury and inflammation. After injury, NO increases expression of cyclooxygenase-2 (COX-2) and enhances COX-2-mediated prostaglandin formation, a well-established pain mechanism [64]. Treating mouse fibroblasts with 7α,25-OHC has been shown to increase COX-2 expression, and ERK signaling boosts inducible NOS (iNOS) expression/activity and inhibits neuronal NOS (nNOS) activity [6; 13; 14; 39]. Both iNOS and nNOS are implicated in models of inflammatory and neuropathic pain, where NOS-induced sensitization is mainly linked to elevated iNOS expression on inflammatory macrophage phenotypes [8; 17; 42; 44; 61]. Moreover, macrophage NO production is a key driver of metabolic reprogramming toward inflammatory functions [53].

Although 7α,25-OHC–mediated GPR183 activation is known to induce MAPK/ERK signaling and increase COX-2 expression, its relationship to NOS activity has not been examined. Here, we investigated whether GPR183 activation promotes peripheral sensitization through ERK-and NOS-dependent mechanisms [14; 25; 36; 56]. In addition, given gaps in our understanding of the cellular contributors to the nociceptive circuit, we examined the source of the predominant GPR183-expressing cells in human incisional injury. We present evidence that GPR183 induces mechanical and thermal hypersensitivities through ERK and NOS, and that this process contributes to peripheral injury-related sensitization. These studies are the first to link NO signaling to GPR183 activity and describe a mechanism by which activation of GPR183,expressed by macrophages, leads to peripheral sensitization.

## 2. Materials and Methods

### 2.1. Animals

Male and female Sprague Dawley rats (200-250g starting weight; 142 total) were purchased from Envigo-Harlan Laboratories (Indianapolis, IN). Age matched (9-11 weeks old) male and female C57BL/6J (WT; Strain #: 000664; 26 total), iNOS KO (Strain #: 002609; 8 total), nNOS KO (Strain #: 002986; 14 total), and Rag1 KO (Strain #: 002216, 4 total) were purchased from Jackson Laboratories (Bar Harbor, ME). Humanized langerin-diphtheria toxin receptor mice (hLangerin-DTR; 14 total) used for LC ablation via diphtheria toxin were maintained as a colony rederived from frozen sperm in the laboratory of Dr. Menichella at the Northwestern Center for Comparative Medicine [51; 82]. Rats were housed 2-4 per cage and mice were housed 4-6 per cage in controlled environments (12-hour light/dark cycle) with food and water available *ad libitum*. Experimental treatment groups were assigned randomly, and rats and mice were housed and tested in separate rooms. All experiments were performed with experimenters blinded to the treatment paradigms. All experiments were performed in accordance with the rules and guidelines from the International Association for the Study of Pain and the National Institutes of Health. The studies were approved by the Saint Louis University Animal Care and Use Committee.

### 2.2. SAE-14 synthesis

SAE-14 was synthesized by first reacting 1,3-difluoro-5-nitrobenzene with morpholine in the presence of K_2_CO_3_ at 120 °C to afford 4-(3-fluoro-5-nitrophenyl)morpholine. 4-(3-Fluoro-5-nitrophenyl)morpholine was then dissolved in methanol, and 10% Pd/C was added and shaken in an atmosphere of 60 PSI hydrogen for 18 hours to yield 3-fluoro-5-morpholinoaniline. 3-Fluoro-5-morpholinoaniline and 3,4-difluorohydrocinnamic acid were stirred in dry DMF in the presence of HATU and triethylamine for 18 hours to yield crude SAE-14. SAE-14 was then purified with reverse-phase column chromatography (Acetonitrile:Water) to a purity of 98%, and was characterized with ^1^H NMR, ^13^C NMR, and HPLC/HRMS.

### 2.3. Test compounds

7α,25-dihydroxycholesterol (cat: 700080) was purchased from Avanti Polar Lipids (Alabaster, AL) and dissolved in DMSO as a 1 mg/mL (2mM) stock. For injections, 7α,25-OHC was serially diluted to 0.1 µg/mL (0.2 µM; rats) or 0.2 µg/mL (0.4 µM; mice) with DMSO and saline (0.9%). Vehicles for 7α,25-OHC were 0.1% DMSO (rats) and 0.2% DMSO (mice). SAE-14 was dissolved fresh from powder in DMSO to a starting concentration of 4 mg/mL (11 mM; 7α,25-OHC injection paradigm) or 2.4 mg/mL (6.6 mM; incision model) and serially diluted with DMSO and saline to 0.52 µg/mL (1.4 µM; 7α,25-OHC injection paradigm) or 12 µg/mL (33 µM; incision model) for injections. Vehicles for SAE-14 contained either 1.3% (7α,25-OHC injection) or 2.5% (incision model) DMSO in saline. U0126 was purchased from Millipore Sigma (St. Louis, MO) and dissolved in DMSO as a 4 mg/mL (10mM) stock and diluted to 0.2 mg/mL (0.5 mM) in saline for injections (5% DMSO in saline vehicle). NS-398 (cat: 70590), SC-560 (cat: 70340), L-NAME (cat: 80210), and L-NIL (cat: 80310) were purchased from Cayman Chemical. NS-398 and SC-560 were freshly reconstituted to 10 mg/mL (3.18 mM and 28.3 mM respectively) stocks in DMSO. NS-398 and SC-560 were diluted to 0.2 mg/mL (0.636 mM and 0.566 mM respectively) in Tween 80, and saline. Vehicles contained 2% DMSO and 1% Tween 80 in saline. L-NAME and L-NIL stocks were dissolved fresh from powder in saline and diluted to 1 mg/mL (3.7 mM) and 0.6 mg/mL (2.3 mM) respectively in saline for injections. ARL 17477 (cat: HY-107720) was purchased from MedChemExpress (Monmouth Junction, NJ); stock solutions were dissolved fresh from powder in saline and diluted to 0.4 mg/mL (0.9 mM) for injections using saline. Diptheria Toxin (DT) was obtained from Sigma-Aldrich (1mg, D0564) and diluted in saline to a final concentration of 4 µg/kg concentration.

### 2.4. Intraplantar injections

Intraplantar injections adapted a previously published protocol [23]. Injections were administered following a baseline behavioral measurement. Rats were anesthetized and maintained on 2.5% isoflurane and 2% O_2_. Hind paws were extended and injected with 50 µL of test agent or its vehicle. Prophylactic drug treatments involved two separate 50 µL injections administered 15-20 minutes apart where the first injection was an antagonist/inhibitor test drug or its vehicle, and the second injection contained 7α,25-OHC or its vehicle. Mice were anesthetized and maintained on 2% isoflurane and 2% O_2_. Mouse hind paws were expressed and injected with 5 µL of 7α,25-OHC or vehicle. No animal behaviors were excluded from these experiments.

### 2.5. Animal behavioral testing

Mechanical and thermal hyperalgesia behaviors were assessed. When measured on the same animal, mechano-allodynia was measured first with at least 30 minutes of habituation before testing thermal hyperalgesia.

#### Mechanical allodynia

was measured as previously described in rats and mice using the Chaplan method [15]. Mechanical allodynia outputs were defined as a significant (*p*<0.05) reduction of mechanical paw withdrawal threshold (PWT) or 50% withdrawal threshold in grams compared to baseline (before treatment or surgery). Rats and mice were placed in cages on an appropriately sized mesh stand and acclimated for at least 30 minutes prior to testing. Calibrated von Frey filaments (Stoelting; range in rats: 1.40-26.00 g; range in mice: 0.017-2.00 g) were applied to the plantar surface of the hind paw two-to-three times with an interstimulus interval of at least 15 seconds where filament contact with hind paw insults (i.e., incision and injection) were avoided (PWT). For LC ablation experiments, mice were habituated for 60-minutes on a metal mesh floor under a transparent plastic dome. The plantar surface of the hind paw of hLang-DTR mice were stimulated with seven filaments with increasing bending force. Each filament was applied six times, with an interstimulus interval of 10–15 s (50% withdrawal threshold) [5; 45; 46; 81].

#### Thermal hyperalgesia

was measured by the Hargreaves test [33]. Rats were placed on a framed glass panel and allowed to acclimate for at least 30 minutes. An infrared heat source was positioned under the hind paw where paw withdrawal latency (PWL) in seconds was recorded with a cutoff of 20 seconds to prevent tissue damage. The test was repeated twice with an interval of 5 minutes between application for each paw. PWL was reported as an average of each test.

### 2.6. Paw incision

The incision model was developed to replicate human post-operative pain in mice and rats [11]. Baseline behaviors were measured prior to surgery. Rats were anesthetized and maintained using 2-3% isoflurane and 2% O_2_. Rats received an intraperitoneal injection of antibiotic, enrofloxacin (Baytril, 3.4 mg). One of the hind paws was sanitized using a chlorohexidine solution and a 1 cm longitudinal incision was made 0.5 cm away from the proximal edge of the heel using a #10 scalpel blade through the skin and fascia directed toward the toes of the hind paw. The plantar muscle tissue was then elevated and cut longitudinally, and sterile gauze was used to apply pressure to the wound. After hemostasis, the skin was apposed with 2 horizontal mattress sutures of silk thread (5-0). Following wound closure, 2% lidocaine gel and triple antibiotic ointment (polymyxin B, neomycin, and bacitracin) were separately applied to the incision site. Behavior was measured 2-3 hours following incision. Sutures were removed under anesthesia (2-3% isoflurane) at the end of postoperative day 1, after mechanical hypersensitivity testing for day 1 and approximately 24-30 hours post-surgery. Sham surgeries involved anesthetizing animals for approximately 5 minutes, administering an intraperitoneal injection of antibiotic, sanitizing paws, and performing a similar cutting and apposition protocols using blunt objects in the same regions of the paw. Lidocaine gel and triple antibiotic ointment were then applied. Approximately 24-30 hours following surgery, sham animals were anesthetized again for approximately 2 minutes to mimic suture removal. Behaviors were then measured daily up to 3 days with increased measurement increments following SAE-14 treatment. Tissues were harvested on day 3 following hind paw incision surgery. Data from animals that removed sutures prior to the within 24-30 hours post-op were excluded from the study (1 animal excluded).

### 2.7. Western blot

Contralateral and ipsilateral hind paw and lumbar dorsal root ganglia (DRG; L4-L6) tissues were harvested and snap frozen from phosphate-buffered saline perfused rats three days following hind paw incision surgery. Positive control spleens were harvested from unperfused WT C57BL/6 mice and snap frozen [30]. Paw tissues were pulverized in a liquid nitrogen-cooled pulverizer and then ground into powder in liquid nitrogen cooled mortar/pestles. The tissues were transferred to microcentrifuge tubes containing 4 volumes of ice-cold homogenization buffer [50 mM Tris-Cl, pH 8.0, 150 mM NaCl, 0.5% Triton X-100, 0.1% SDS, 1 mM EDTA, 5% glycerol, 1 mM PMSF, 1X protease inhibitor cocktail (#P8643, Millipore-Sigma), 1 mM Na_3_VO_4_, 1 mM Na_3_MbO_4_, 50 mM NaF and 1X phosphatase inhibitor cocktail (#P0444, Millipore-Sigma)] and pulse sonicated (Q800R Sonicator Chromatin and DNA Shearing System, Qsonica; Newtown, CT) in a 4⁰C cold water bath at 90% amplitude for 5 minutes (30 seconds on, 45 seconds off). DRGs and spleens were first homogenized in tubes submerged in ice using microcentrifuge pestles in 10-12 volumes (DRG) or 4 volumes (spleen) of ice-cold buffer and subsequently pulse sonicated at 85% amplitude for 5 minutes (30 seconds on, 45 seconds off) in a 4⁰C cold water bath. The samples were clarified by centrifugation at 15,000 x g, 15 minutes at 4°C. The total protein concentration in the clarified lysates was measured using bicinchoninic acid (Thermo Fisher Scientific, Carlsbad CA, USA). Lysate proteins (50 µg for paws and 20 µg for DRG) were resolved by TGX stain-free 10% sodium dodecyl sulphate-polyacrylamide gel (BioRad, #5678034; Hercules, CA) electrophoresis, stain-free activated (5-minute activation) and transferred to low fluorescence polyvinylidene difluoride (PVDF) membranes (0.45 µM pore size). Membranes were blocked for 1 hour at room temperature in a solution containing 3% bovine serum albumin (#A7030, Millipore-Sigma) and 0.01% thimerosal (#T5125; Millipore-Sigma) in 1X tris-buffered saline (TBS; 15 mM Tris pH 7.6, 150 mM NaCl) and cut just below the 37 kDa molecular weight. The top of the blot (>37 kDa) was incubated with antibodies specific to GPR183 (Invitrogen, PA1-032; 1:500-1000) in 1X TBS containing 1% BSA and 0.003% thimerosal at 4°C overnight. The membranes were washed in 1X TBS-Tween-20 (TBS-T; 15 mM Tris pH 7.6, 150 mM NaCl, 1% Tween-20) for three times for 10 minutes per wash and incubated with horseradish peroxidase (HRP)-conjugated goat anti-rabbit IgG (Cell Signaling Technologies, 7074S; 1:5000) for 1 hour at room temperature. The blot was washed again three times for 10 minutes per wash with 1X TBS-T, and labeled proteins were visualized by enhanced chemiluminescence (Clarity or West Femto, Bio-Rad; Hercules, CA) and documented using Chemidoc MP and ImageLab software (BioRad). Blots were treated for fifteen minutes with 30% hydrogen peroxide to deactivate the HRPO to allow visualization of a second target [69]. The membrane was then washed for 15 minutes total (three times for 5 minutes per wash) with TBS-T and treated with antibodies targeting α-tubulin (Millipore Sigma, T8203; 1:2000) at 4°C overnight. The membranes were washed in 1X TBS-T three times for 10 minutes per wash. The bound antibodies were then visualized following incubation with HRP-conjugated goat anti-mouse IgG secondary antibody (SeraCare, 5220-0287; 1:5000) for 1 hour at room temperature. GPR183 and α-tubulin bands were visualized by enhanced chemiluminescence (Clarity or West Femto, Bio-Rad; Hercules, CA) and documented using Chemidoc MP and ImageLab software (BioRad). Images were selected based on a histogram maximum of 30,000-35,000 (GPR183) and 40,000-45,000 (α-tubulin). Relative GPR183 expression in each lane was quantified by measuring band densitometry and normalized to the α-tubulin density of that lane from the same blot using the lane and band functions of the ImageLabTM software. For publication, the image contrast was set to 30000 high and 0 low, gamma was set to 1.00 in ImageLabTM prior to exporting as.tiff files. All blots were run with spleen lysate as a positive control for GPR183 [30]. Samples lysates (36 total tissues) were analyzed on three separate blots and repeated analysis showed similar results.

### 2.8. Bioinformatic analysis

Transcriptomics expression data and significance values from rat surgical incision model was mined from a previous publication [31]. In this dataset, transcriptomics were performed on excised hind paw samples collected in the midplantar region, including epidermis, dermis and muscle following surgical injury. To extend these findings to human subjects, an additional analysis was performed based on data collected as part of a registered clinical trial (NCT04224870). A supplementary analysis was performed using data from a previously published database of transcriptomic studies examining rat DRG responses to nerve injury [65].

### 2.9. Human tissue samples

Human skin samples were collected during surgery under an approved protocol at the National Institutes of Health (Bethesda, Maryland; 20-CC-0031) and the protocol was registered at ClinicalTrials.gov (“A single cohort study collecting interval timed incisional epidermal and dermal tissue samples during surgical procedures to profile temporal response of tissue after noxious stimuli;” NCT04224870). Informed consent was obtained from all participants prior to enrollment in the study.

### 2.10. Multiplex fluorescence in situ hybridization and microscopic image capture

Multiplex in situ hybridization of human skin samples (N=3) was performed using the RNAScope® Version 2 kit (Advanced Cell Diagnostics, Newark, CA) following a modified procedure as described previously [66], and visualized using Opal™ Reagent Systems (Akoya, Marlborough, MA). Target retrieval was performed for ∼20 minutes at 100°C. Digestion was performed using the Protease Plus reagent. Image capture of hybridized stained slides was performed using an Axio Imager. Z2 scanning fluorescence microscope (Zeiss, Oberkochen, Germany) as described previously [31]. For final visualizations, single-channel microscopy images captured at each emission wavelength were overlaid to generate multi-colored composites in Adobe Photoshop (San Jose, CA). Representative images are enhanced for visibility.

### 2.11. Diphtheria toxin-mediated ablation

To deplete LCs, hLangerin-DTR mice received weekly injections of Diphtheria toxin (DT; 4 µg/kg) from 10 to 14 weeks of age. Control mice received DT vehicle: saline (0.9% NaCl). Approximately 48h following final DT or saline injection, behavioral assessments were performed and paw epidermis was harvested. Efficiency of LC ablation was assessed by quantifying Cd207^+^ LCs in epidermal whole-mount preparation. Two animal’s withdraw thresholds were calculated as greater than our maximum filament (2 g); therefore, these animals were excluded from study as baseline measurement was out of range.

### 2.12. Whole-mount staining

To visualize LCs in the mouse paw skin, a whole mount staining of the epidermis was performed as previously described [51]. Briefly, glabrous hind paw skin was incubated in Dispase (2.3 mg/ml, Sigma-Aldrich D4693) overnight at 4C. The following day, the epidermis was separated from the dermis and the epidermal tissue was fixed in 4% PFA for 30 minutes at RT. The tissue was permeabilized with 0.2% Triton X-100 in PBS, then incubated in a blocking solution containing 10% NDS - 0.1% BSA in 0.1% Triton X-100-PBS for 1hour at RT and incubated with APC anti-mouse/human Cd207 1:100 (Biolegend, 4C7) overnight at 4C. Images were acquired using a confocal fluorescence microscope.

### 2.13. Statistical analysis

All data collected were analyzed and expressed as mean ± standard deviation. Behavior data was analyzed using two-tailed, two-way ANOVA with Bonferroni *post hoc* comparisons. Protein expression and LC density data was interpreted using two-tailed, unpaired t-test. All data were analyzed by using GraphPad Prism (v10.2.3; GraphPad, San Diego, CA). All bioinformatic graphs were constructed in Prism 10 (GraphPad, Boston MA). Significant differences were defined at *p* < 0.05. For RNA-Seq analyses, statistical methods where differential genes were identified using the MAGIC pipeline for longitudinal data from rat and human skin incision as reported in original publications [31; 67].

## 3. Results

### 3.1. Peripheral GPR183 induced behavioral hypersensitivities through MAPK

Intraplantar injection of 7α,25-OHC in male and female rats evoked time-dependent and long-lasting development of mechano-allodynia that persisted for up to 5 days; these effects were blocked by the selective GPR183 antagonist, SAE-14 (**Fig. 1A**). In addition, intraplantar injections of 7α,25-OHC induced thermal hyperalgesia that began ∼1h following injection of 7α,25-OHC and resolved within 24 hours, and this was prevented by SAE-14 (**Fig. 1B**). No changes were observed in contralateral paws (**Supplemental Fig. 1A-B**). Our previous studies linked ERK signaling downstream of GPR183 in the spinal cord that contributed to central sensitization [9]. To determine if a similar pathway was engaged in the paw tissue for the development of peripheral sensitization, we tested the effects of an intraplantar injection of the MEK 1/2 inhibitor, U0126 [24; 83]. Intraplantar injection of U0126 blocked the development of 7α,25-OHC-induced mechanical allodynia (**Fig. 1C**) and thermal hyperalgesia (**Fig. 1D**) induced by 7α,24-OHC. No changes were observed in contralateral paws (**Supplemental Fig. 1C-D**).

**Figure 1:**
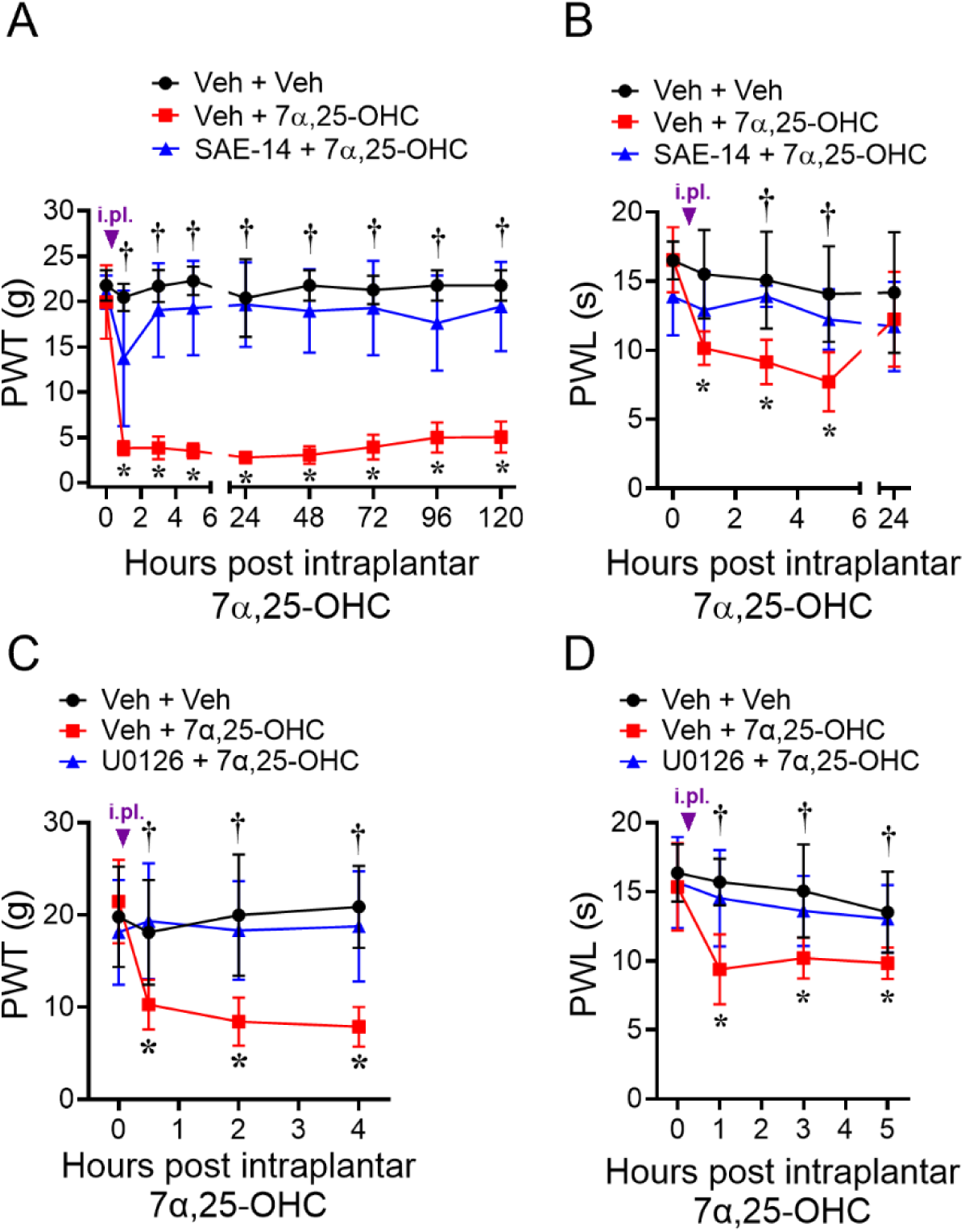
Intraplantar injection of 7*α*,25-OHC induces sensitization in rats through MAPK/ERK. **A,B)** Intraplantar injections of 7α,25-OHC (5 ng) induce time-dependent development of mechanical allodynia **(A)** and thermal hyperalgesia **(B)** in male and female rats that are blocked by pretreatment with a selective GPR183 antagonist, SAE-14. **C,D)** 7α,25-OHC-induced (5 ng) mechanical allodynia **(C)** and thermal hyperalgesia **(D)** were blocked by pretreatment a selective MEK1/2 inhibitor (U0126, 10 µg). Data are expressed as mean ± SD for n=6/group **(A,C,D)** or n=4/group **(B)** and analyzed by two-tailed, two-Way ANOVA with Bonferroni multiple comparisons: \**p*<0.05 vs 0 hours; †*p*<0.05 vs. Veh+7*α*,25-OHC group.

### 3.2. Nitric oxide contributes to the development of GPR183 signaling

Prior studies have identified a link between ERK and nitric oxide synthases [6; 13; 14], which, coupled with the known involvement of NOS in pain states [23; 42; 62; 64], raised the intriguing possibility that NOS activity may contribute to GPR183-mediated effects. This was tested using three well-characterized NOS inhibitors given by intraplantar injections in rats; L-NAME, a non-selective NOS inhibitor [52], L-NIL, an iNOS selective inhibitor [48], and ARL 17477, an nNOS selective inhibitor [84]. Doses were chosen from previous studies [12; 72; 74]. L-NAME attenuated the development of 7α,25-OHC-induced mechano-allodynia and accelerated recovery of baseline values by 48 hours (**Fig. 2A**). L-NIL partially attenuated mechano-allodynia but did not accelerate recovery (**Fig. 2B**) whereas ARL 17477 did not attenuate mechano-allodynia but began to accelerate recovery after 4 hours with a complete return to baseline by 96 hours (**Fig. 2C**). These effects were confirmed in iNOS and nNOS knockout mice (**Fig. 2D and E**) and there were no changes observed in the contralateral paws of rats or mice (**Supplemental Fig. 2A - E**). Nitric oxide is known to increase the expression and activity of COX-2 which catalyzes the formation of prostaglandins from arachidonic acid and exacerbates pain symptoms [64]. We hypothesized that GPR183 activation would result in increased COX-2 activity that impacts hypersensitivities. We selectively blocked COX-1 and COX-2 activity using intraplantar injections of SC-560 [54] and NS 398 [28] respectively approximately 15 minutes prior to injection of 7α,25-OHC. Doses for these inhibitors were chosen from a previous study [68]. We found that inhibition of COX isoforms had no effect on 7α,25-OHC-induced mechano-allodynia, suggesting that the GPR183-mediated NOS mechanism that induces hypersensitivities is independent of COX (**Fig. 2F**). No changes were observed in the contralateral paws (**Supplemental Fig. 2F**).

**Figure 2:**
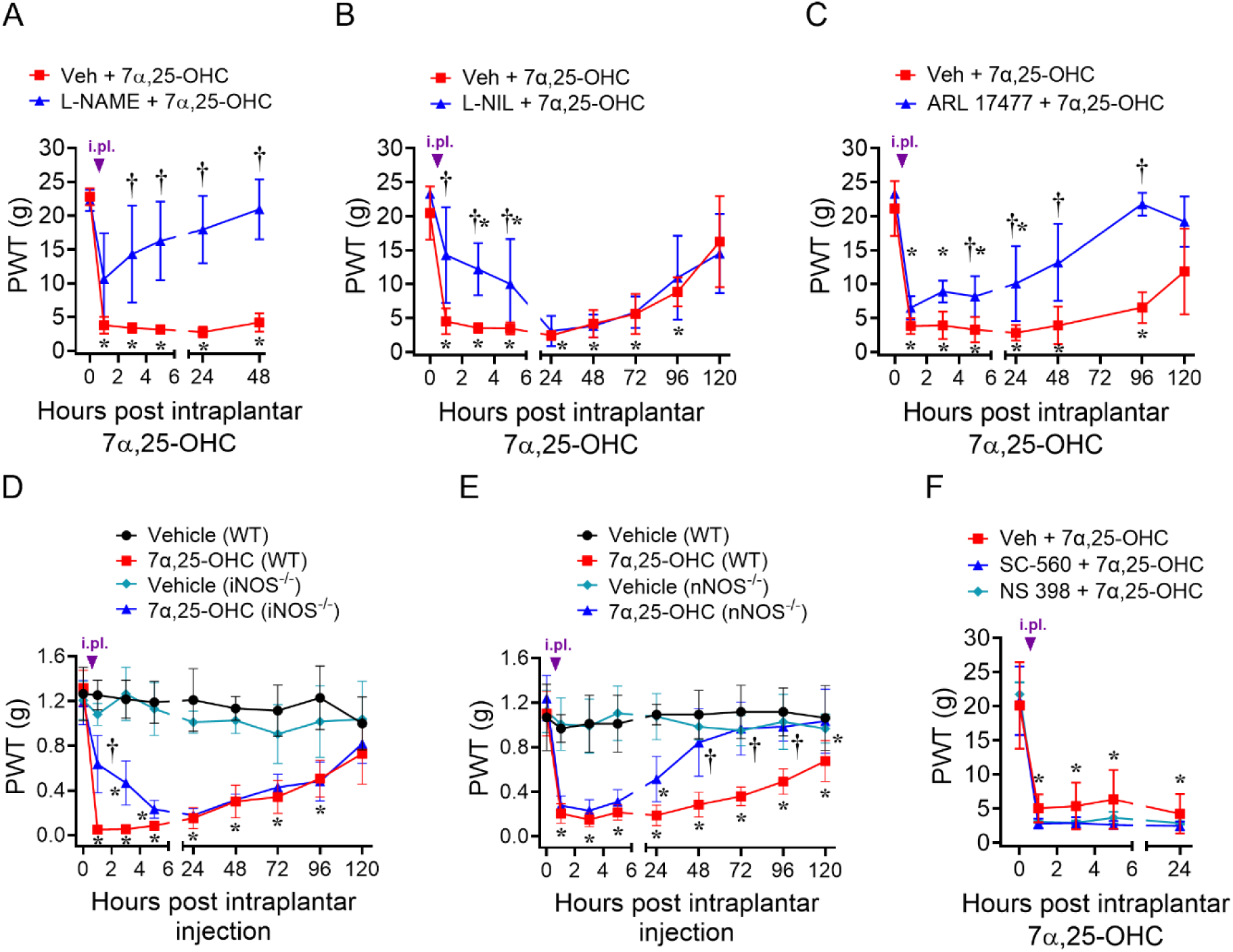
Allodynic effects of 7*α*,25-OHC are dependent on NOS but not COX. **A)** Mechanical allodynia from intraplantar injection of 7*α*,25-OHC (5 ng) was attenuated acutely and prevented long-term in male rats by pretreatment with a non-selective NOS inhibitor (L-NAME, 50 µg). **B)** Pretreatment with a selective iNOS inhibitor (L-NIL, 30 µg) prevented acute mechanical allodynia with no effect on persistence. **C)** Pretreatment with a selective nNOS inhibitor (ARL 17477, 20 µg) accelerated baseline recovery with mild effects on acute hypersensitivity. **D, E)** NOS inhibitor effects were recapitulated when iNOS KO (iNOS-/-) **(D)** and nNOS KO (nNOS-/-) **(E)** mice received an intraplantar injection of 7α,25-OHC (1 ng). **F)** Pretreatment with intraplantar an COX-1 inhibitor (SC-560, 10 µg) and COX-2 inhibitor (NS-398, 10 µg) in male rats had no effect on 7α,25-OHC-(5 ng) induced mechanical allodynia. Data are expressed as mean ± SD for n=6/group **(A,B,C),** n=4/group **(D,F)**, or n=7/group **(E)** and analyzed by two-tailed, two-Way ANOVA with Bonferroni multiple comparisons: \**p*<0.05 vs 0 hours; †*p*<0.05 vs. Veh+7*α*,25-OHC group.

### 3.3. GPR183 antagonists reverse paw incisional injury induced hypersensitivities

To further explore the role for GPR183 in peripheral sensitization we used a model of acute post-surgical induced peripheral sensitization in rats (incision model). The incision model is a deep cut to the hind paw in mice and rats used to mimic post-operative pain. The model induces intense pain that gradually heals and returns to baseline after several days [11]. Analysis of a publicly available transcriptomic data set to examine *Gpr183* expression changes in the rat incisional model [31] revealed that *Gpr183* transcription increased in rats steadily with the highest expression reaching approximately 3-fold, three days following incisional surgery (**Fig. 3A**). 7α,25-OHC is synthesized from cholesterol in vivo through two enzyme-catalyzed hydroxylation reactions at the 25 and 7α positions via 25-cholesterol hydroxylase (CH25H) and cytochrome P450 family 7 subfamily B member 1 (CYP7B1), respectively, and is metabolized by hydroxy-delta-5-steroid dehydrogenase, 3 beta-and steroid delta-isomerase 7 (HSD3B7) (**Fig. 3A**) [80]. Constitutive (baseline) expression of 7α,25-OHC synthesis enzymes, *Ch25h* and *Cyp7b1*, are low in rat dermal tissue while expression of the 7α,25-OHC degradative enzyme, *Hsd3b7*, is relatively high (**Fig. 3A, Supplemental Fig. 3**). Expression of *Ch25h* is induced one hour following injury, *Cyp7b1* expression increases starting around six hours through three days, and expression of *Hsd3b7* decreases from for the duration of one hour to 24 hours with increased expression at 6 days; a shift of expression of these enzymes that aligns with expected *Gpr183* expression changes (**Fig. 3A**). These results suggest a potential role for GPR183 in peripheral injury leading to an independent investigation in the rat model of incisional injury (**Fig. 3B**). GPR183 protein expression was significantly elevated in hind paws receiving surgery compared to their non-injury side three days after injury (**Fig. 3C, Supplemental Fig. 4A**). This increase in GPR183 expression did not extend to the DRG tissues at this time point (**Supplemental Fig. 4B-C**). Paw incision resulted in severe mechanical allodynia that was reversed approximately 50-60% following intraplantar injection of SAE14 (**Fig. 3D**). No changes were observed in the contralateral paws (**Supplemental Fig. 4D**).

**Figure 3.**
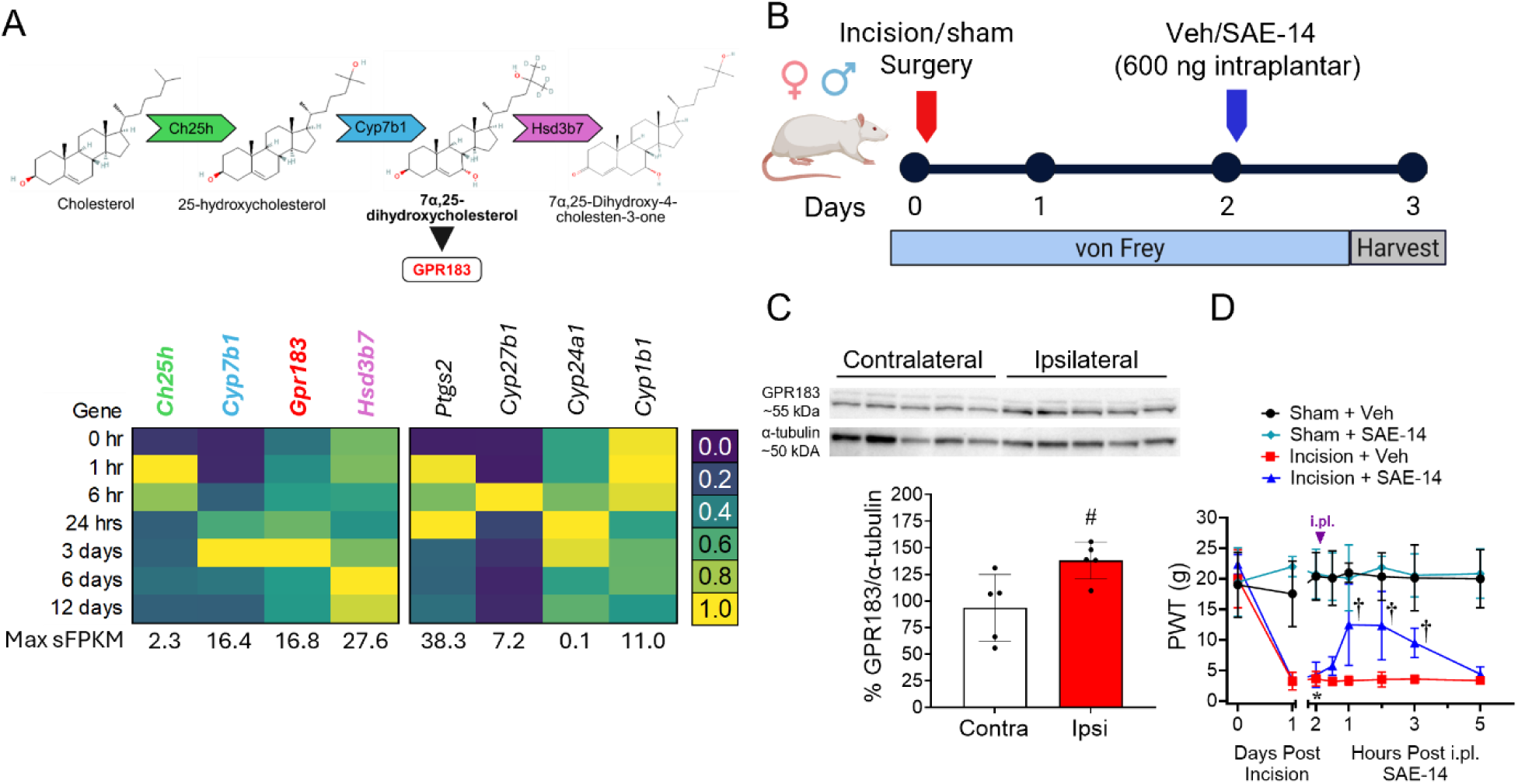
GPR183 expression is induced after peripheral injury in rat dermal tissue; GPR183 antagonists reverse behavioral hypersensitivity. **A)** A publicly available rat paw incision transcriptomic dataset [31] was mined and showed that *Gpr183* was significantly induced, and that expression of 7α,25-OHC synthesis and degradation enzymes *(Ch25h, Cyp7b1, Hsd3b7)* shifted toward 7α,25-OHC synthesis. **B)** Incision model timeline for behavior and tissue harvest in male and female rats. **C)** GPR183 protein expression was elevated in incised paw tissues compared to naïve paw with repeated experiments having similar results. **D)** Intraplantar injection of GPR183 antagonist (SAE-14, 600 ng) partially reversed incisional injury pain. Data are expressed as mean ± SD for n=5/group **(C)** or n=7/group **(D)** and analyzed by **(C)** two-tailed, unpaired t-test, #*p*<0.05 vs Contra, or **(D)** two-tailed, two-way ANOVA with Bonferroni comparisons: \**p*<0.05 vs. 0 hours; †*p*<0.05 vs. Incision + Veh.

### 3.4. *GPR183* expression in human skin tissue over the course of long surgeries

To confirm that transcriptional changes observed in rats (**Fig. 3A**) [31] also occur in humans, we examined both basal expression values of *GPR183* in human RNA expression databases, and expression changes in *GPR183* after surgical incision in humans.

Using publicly available datasets, *GPR183* expression is consistent with a role in the innate immune system with an expression pattern consistent with functions in lymphoid and bone marrow [60; 75]. As the present study chiefly examined peripheral functions of GPR183 in the skin, specific single-cell analyses were performed to predict *GPR183* expression patterns in human skin cells. This analysis revealed that *GPR183* expression was enriched in dermal macrophages/dendritic cells and T-cells in human skin (**Supplemental Fig. 5**) [73]. Moreover, review of the primary data from two other independent single-cell RNA-Seq datasets confirm expression of *GPR183* in LCs and macrophages [38; 60; 75].

Similar to the rat hind paw, *GPR183* was induced strongly in human skin during surgery, continually increasing throughout the surgical case. This induction was notably faster than circulating macrophage recruitment (**Fig. 4A**) [67], indicating that it is most likely due to gene regulation by localized immune cells (e.g., resident macrophages, T cells, and LCs), and not recruitment of circulating macrophages to the wound edge [41]. Constitutive (baseline) expression was detectable in human skin for *CH25H* and *CYP7B1*, although expression (significant fragments per kilobase per million aligned reads, sFPKM) for *CYP7B1* was low throughout the study. Additionally, both enzymes were induced at 6 hours in rat and human, consistent with *GPR183* expression peaking at 6 hours after injury, which is the peak sensitization point after tissue injury and/or inflammation [78]. The *HSD3B7* transcript is expressed at high levels in the nociceptive circuit (**Supplemental Fig. 3**) but is not consistently regulated by injury in rats and humans (**Figs. 3A and 4A**). Overall, these data establish that expressions of *GPR183* and synthesis enzymes of 7α,25-OHC are significantly induced by tissue injury (**Fig. 4A**).

**Figure 4.**
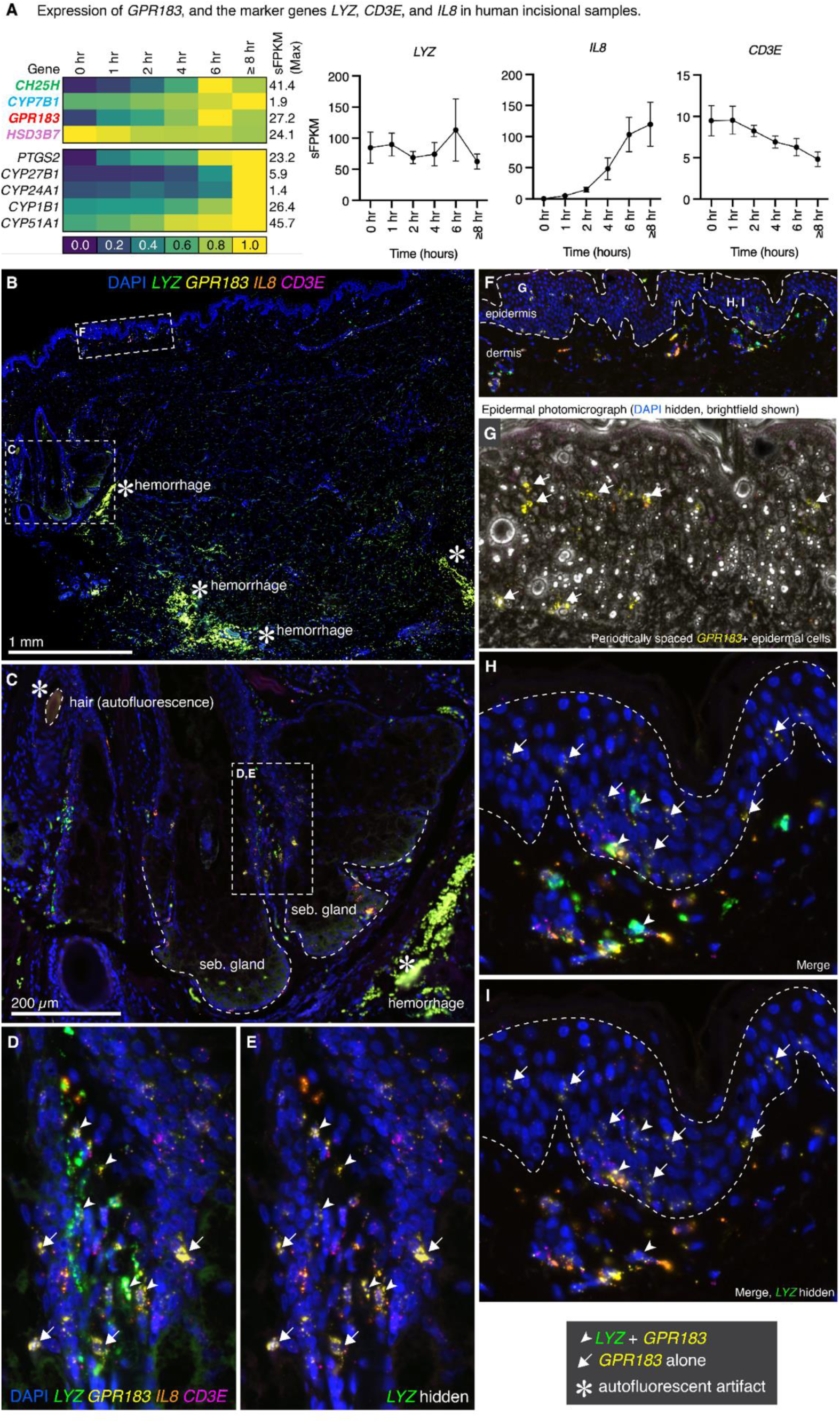
Epidermal and dermal spatial expression of *GPR183* in human skin samples. **A)** Graphs showing the expression of *GPR183*, *LYZ*, *IL8*, *CD3E* across time in a human incision model. *GPR183* was significantly induced after human surgical incision. Of note is *GPR183* induction in context of relatively stable *LYZ* expression. **B-I**) Human skin samples were bed scanned from large slabs that were serially collected from a wound edge during long surgical cases (see Methods) and show positive staining for GPR183. **B)** Full visualization of skin slab. Severe hemorrhage appeared as highly autofluorescent red blood cells at the wound edge (asterisks), indicating tissue damage 6 hours into surgery. Spatial analysis was performed to examine the anatomical subregions showing the highest *GPR183* signal. **C)** Magnification of sebaceous glands with abundant *GPR183* expression (note the autofluorescent crossection of a hair in the upper-left marked by asterisk). **D, E)** Enlarged view of the interface between two sebaceous gland lobes shows an accumulation of **(D)** *LYZ*^+^/*GPR183*^+^ secretory macrophages (arrowheads) and **(D, E)** *GPR183*^+^ immune cells only (arrows). **F)** Magnification of the epidermis shows *GPR183* signal throughout epidermal and subepidermal regions. **G)** To examine putative LCs, we visualized a subregion of epidermis marked by single-labeled *GPR183*^+^ cells evenly spaced throughout the epidermis, consistent with LCs positive for this receptor. **H, I)** Enlargement of epidermal/dermal interface, **(H)** *GPR183*^+^/*LYZ*^+^ secretory macrophages and **(H, I)** GPR183^+^ cells were visible at specific subregions of the tissue. In all panels *CD3E* showed limited co-expression with *GPR183*, suggesting limited contribution of *GPR183* expression from T-cells in human skin.

Based on the predicted cell type expression from single-cell RNA-Seq of human skin, a 4-plex probe set was designed to specifically interrogate cellular expression of *GPR183* in human skin collected during surgery (**Fig. 4B-I**). Large skin tissue slabs were collected during surgery and probed for human *GPR183* with additional markers for T-cells (*CD3E*) and macrophages (*LYZ*) in human skin [73] alongside interleukin 8 (*IL8*), which is one of the most highly significant induced genes in human skin in macrophage-like cells [67]. Overview of the entire tissue reveals expression of these markers associated with various dermal structures with some regions of highly autofluorescent red blood cells due to hemorrhage at wound edge (**Fig. 4B**). Notable structures with abundant GPR183 signal are the sebaceous glands (**Fig. 4C-E**) and the epidermis (**Fig. 4F-I**). Closer inspection of the sebaceous glands reveals an accumulation of *LYZ*^+^/*GPR183*^+^ and *GPR183*^+^ only cells surrounding this structure (**Fig. 4C**) particularly in the region between the sebaceous gland lobes (**Fig. 4D-E**). Magnification of the epidermis and subepidermal layer shows *GPR183* expression interspaced throughout this region (**Fig. 4F**). *GPR183* was expressed evenly throughout the epidermal tissue consistent with LC localization patterns (**Fig. 4G**). Further magnification of the epidermal and subepidermal regions show *GPR183* expression with and without association of *LYZ* expression (**Fig. 4H-I**). *GPR183* expression is, again, shown to be evenly spaced throughout the epidermal tissue suggesting expression by LCs, and *GPR183* was also sparsely associated with *LYZ^+^* cells indicating that these cells are either epidermis-resident macrophages and/or *LYZ^+^* LCs (**Fig. 4H-I**). *IL-8* expression was increased in surgical injury and visualized throughout the tissue where it was occasionally associated with *GPR183*^+^ cells (**Fig. 4**). *CD3E* was not strongly associated with *GPR183*, indicating that T cells are not likely involved in GPR183-mediated peripheral surgical pain (**Fig. 4, Supplemental Fig. 6A-C**).

### 3.5. 7*α*,25-OHC-mediated hypersensitivities develop independent of Langerhans cells, T cells, and B cells

In both our human skin tissue and in published transcriptomic datasets in humans, rats, and mice, GPR183 is expressed in both macrophages and LCs, with additional evidence for expression in other immune cell types as addressed above [37; 38; 51; 60; 75]. Given the relative promiscuity of the receptor, we sought to resolve the cell-specific basis of GPR183 induced nociceptive sensitivity. To determine whether LCs contributed to the 7α,25-OHC-induced nociception, we assessed the mechanical sensitivity in hLangerin-DTR mice following DT-mediated LC ablation(**Fig. 5A**). Intraplantar injection of 7α,25-OHC (1 ng) induced robust mechanical allodynia in both DT-ablated and vehicle-treated mice (**Fig. 5B**), suggesting that loss of LCs did not alter the pro-nociceptive effect of 7α,25-OHC. Effective ablation of LCs was confirmed by quantification of Cd207^+^ (langerin) cells in whole-mount epidermis which showed significant reduction in LC density in DT-ablated mice (**Fig. 5C**).

**Figure 5.**
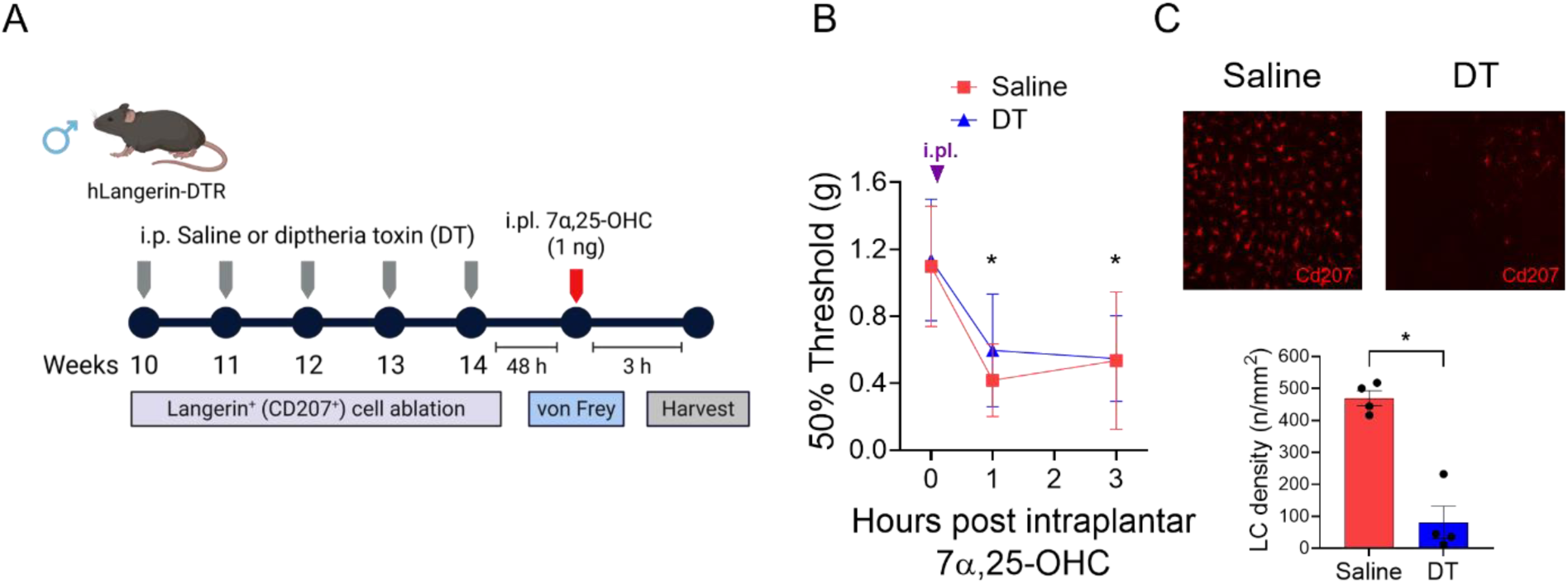
Ablation of Langerhans cells did not prevent the development of 7α,25-OHC-mediated hypersensitivities. **A)** LC ablation model timeline for DT-mediated depletion, behavior, and epidermal tissue harvest in male hLangerin-DTR mice. **B)** Intraplantar injection of 7α,25-OHC (1 ng) induced mechanical allodynia in both LC ablated (DT treated) and control (saline treated) hLangerin-DTR mice. **C)** LC ablation was confirmed in whole mount epidermal Cd207 stains of DT-treated mice. LC density was measured as the number of Cd207^+^ cells/areamm2. Data are expressed as mean ± SD for n=6-8/group **(B)** or n=4/group **(C)** analyzed by **(B)** two-tailed, two-Way ANOVA with Bonferroni multiple comparisons: *p<0.05 vs 0 hours; †p<0.05 vs. Saline group or **(C)** unpaired, two-tailed t-Test with Welch’s correction *p=0.0018.

GPR183 has also been identified on T cells and B cells as an important receptor for migration and positioning [27; 29; 43]. We showed that *GPR183* expression in the human skin following surgical incision does not associate strongly with the T cell marker *CD3E* (**Supplemental Fig. 6A-C**). To determine if 7α,25-OHC-mediated hypersensitivity is dependent on B cells and/or T cells we gave an intraplantar injection of 7α,25-OHC (1 ng) to Rag1 KO mice (**Supplemental Fig. 6D-E**). Rag1 is an essential gene in the maturation of T cells and B cells; therefore, the Rag1 KO mice are deficient in both of these cell types [47]. Both Rag1 KO mice and their age-matched C57/BL6J (WT) counterparts developed mechanical hypersensitivity following an intraplantar injection of 7α,25-OHC suggesting that B cells and T cells do not drive the 7α,25-OHC/GPR183-induced hypersensitivities (**Supplemental Fig. 6D**). No changes were observed in the contralateral paws of either group (**Supplemental Fig. 6E**).

## 4. Discussion

In this study, we identified a GPR183-driven mechanism capable of initiating peripheral sensitization across multiple sensory modalities that was persistent for up to five days (cartoon, **Fig. 6**). Further, *GPR183* shows marked induction in rat and human tissue damage models, demonstrating that control of this transcript is involved in responding to tissue injury. Beyond this, the endogenous machinery for oxysterol production, particularly the enzymes responsible for generating the high-potency agonist 7α,25-OHC, are also upregulated in response to tissue injury, implicating both the ligand and its receptor as “early responders” to acute injury. Furthermore, we provide new evidence for the downstream signaling of 7α,25-OHC-induced GPR183 activation, where we demonstrate that its effects are dependent on MAPK/ERK signaling and activation of NOS in the surrounding tissue. By examining the major cells in human skin in which *GPR183* is induced, we observed *LYZ*^+^ secretory macrophages as major responders to tissue injury, suggesting that GPR183’s actions in the context of tissue damage are largely in this population of cells. Taken together, these findings suggest that early responses to tissue injury mobilize secretory macrophages via oxysterol production and GPR183 signaling. This process likely drives tissue-resident macrophage recruitment [32; 57] and regulates local metabolic states through iNOS, promoting a proinflammatory phenotype [53]. This work provides a foundational mechanistic framework for understanding how the neural–immune interface drives nociceptive sensitization in the skin following tissue injury.

**Figure 6.**
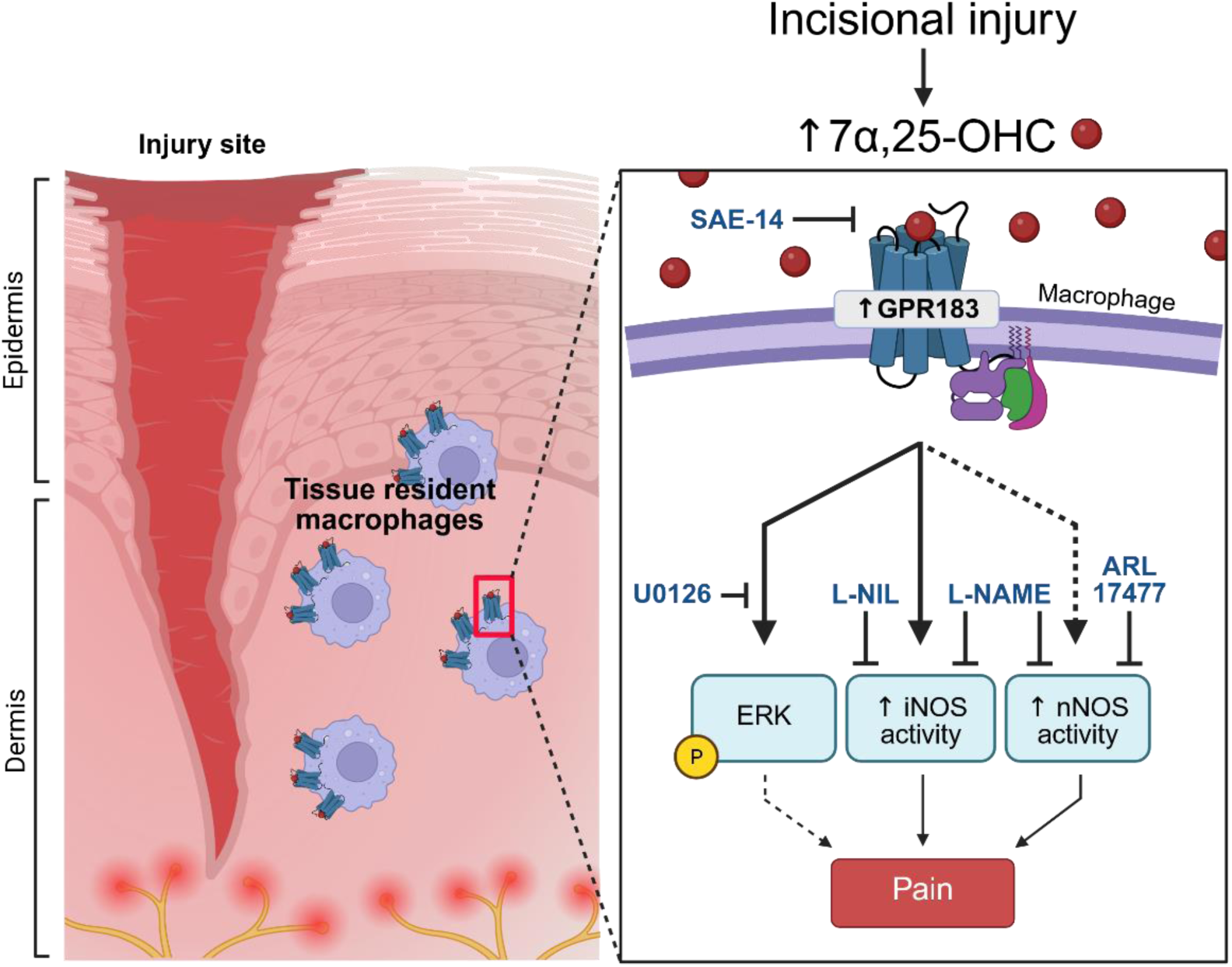
Hypothesized dermal mechanisms of GPR183 activation following incisional injury. Following injury, GPR183 expression is increased in the skin tissue likely through increased expression or chemotaxis on tissue-resident secretory macrophages. GPR183 activated by 7α,25-OHC signals through ERK and iNOS, and downstream from these pathways, constitutively expressed nNOS is activated and maintains pain state persistence.

MAPK/ERK proteins are one of three families of intracellular signaling proteins that transmit a rapid, wide range of transcriptional and non-transcriptional changes in response to extracellular stimuli and is involved in both peripheral and central mechanisms underlying acute and chronic pain conditions [36; 59]. In immune cells, activation of ERK downstream of GPCR signaling induces the release of various inflammatory factors in response to stimuli such as pathogens and injury [49]. Indeed, ERK 1/2 phosphorylation induces activity of NF-κB transcription factors and subsequent release of cytokines, chemokines, and lipid mediators [49], which is a plausible action for GPR183 activation in secretory macrophages that we hypothesize to be involved. Inflammatory mediators released during immune responses also activate ERK pathways in nociceptors that translate into peripheral neuronal sensitization [59]. GPR183 activates ERK [4], and we have found that blocking ERK using local, prophylactic treatment with U0126 was sufficient to prevent the development of peripheral 7α,25-OHC/GPR183-induced hypersensitivities, indicating that downstream ERK signaling is likely required for sensitization of the nociceptive afferents after activation of the GPR183^+^ macrophages by 7α,25-OHC. Importantly, iNOS is amongst the various inflammatory factors induced through ERK/NF-κB signal transduction and transcription during inflammatory responses [49].

NO and NOS have mixed results regarding its role as pro-vs. anti-nociceptive activity [20]. Increased expression and activity of iNOS is strongly linked to hyperalgesia through central and peripheral sensitization, consistent with our results [19]. Similar to work in neuropathic pain models where nNOS inhibition in the spinal cord reversed behavioral hypersensitivities [61; 85], we show that treatment with the selective nNOS inhibitor, ARL 17477, resulted in anti-nociceptive behaviors. While animals given prophylactic ARL 17477 exhibited mild-to-no effects shortly after 7α,25-OHC injection, we observed that recovery of baseline sensitivity was accelerated following ARL 17477 treatment, suggesting that nNOS is active in later stages of GPR183 signaling, where it has pro-nociceptive functions. These findings demonstrate that NO and NOS possess nociceptive functionality and contribute to sensitization processes downstream of peripheral GPR183 activation.

A key contribution of this study is the identification of potential cellular sources of GPR183-induced nociception. While GPR183 is expressed by diverse immune cell populations in mice, rats, and humans [3; 26; 37; 50], we show that its expression is markedly induced after surgical injuries in both rats and humans. Local and selective inhibition of GPR183 activity in rats partially reversed nociceptive sensitivities, implicating this receptor in injury-induced hyperalgesia and allodynia. Importantly, our data point to the *GPR183*^+^/*LYZ*^+^ perifollicular macrophage in human skin as the most likely cellular target of GPR183 antagonists. This secretory macrophage population can be rapidly mobilized by inflammation or tissue damage, releasing pronociceptive mediators such as iNOS and C-C motif chemokine 22, the latter linked to GPR183 signaling in inflammatory pain models [1; 58]. Furthermore, GPR183-mediated chemotaxis of *LYZ*^+^ macrophages may itself promote peripheral sensitization, as lysozyme increases Aδ-and C-fiber excitability and produces persistent pain-like states consistent with our findings following 7α,25-OHC injection [76]. GPR183 expression on macrophages has also been observed in other inflammatory settings, including lung infections [26], and in skin after formalin, CFA, or 7α,25-OHC challenge [34; 58; 77]. LCs were another potential cell target of GPR183 activity as *Gpr183/GPR183* has high expression in this cell type, cross-species, and has been linked to neuropathic pain [21; 37; 38; 60; 71]. Depletion of LCs using DT-treated hLangerin-DTR mice did not prevent 7α,25-OHC/GPR183-induced nociceptive hypersensitivity, suggesting that the LC population is likely not contributing to GPR183-mediated sensitivities. Additionally, T cells were a potential cell target as they have been linked to GPR183 mechanisms and have well known involvement in pain [22; 27]. We found that T cells did not strongly express GPR183 in human skin and that deficiency of mature cells did not have an effect on 7α,25-OHC/GPR183-mediated pain, matching a finding where skin resident T cells, γδ T cells, modulated the immune cell recruitment, but did not contribute to pain during peripheral inflammation [55]. Future studies incorporating additional loss-of-function strategies, such as DTR–mediated, macrophage-specific depletion, will more definitively establish the contribution of macrophages to GPR183 signaling. Collectively, these findings indicate that GPR183 is dynamically induced by tissue injury, contributes to nociceptive signaling—most likely through macrophage-dependent secretory mechanisms—and represents a promising target for the development of novel pain therapeutics.

## Acknowledgements

The authors thank Dr. Daniel Kaplan for kindly providing the hLangerin-DTR mouse sperm for colony establishment. We would like to thank Dr. Timothy M. Doyle for editorial and statistical assistance. We kindly thank Israel Olayide for assistance with synthesizing SAE-14. We would also like to thank Zhoumou Chen and Dr. Luigi A. Giancotti for their surgical, harvesting, and behavioral assessment support. Visual abstract, 7α,25-OHC synthesis pathway, and experimental timelines were created with Biorender.com. The 7α,25-OHC synthesis pathway image uses images of Cholesterol (CID 5997 https://pubchem.ncbi.nlm.nih.gov/compound/5997#section=2D-Structure), 25-hydroxycholesterol (CID 65094 https://pubchem.ncbi.nlm.nih.gov/compound/65094#section=2D-Structure), 7α,25-OHC (CID 11954197 https://pubchem.ncbi.nlm.nih.gov/compound/11954197#section=2D-Structure), and 7α,25-dihydroxy-4-cholesten-3-one (CID 5284267 https://pubchem.ncbi.nlm.nih.gov/compound/5284267#section=2D-Structure). These studies were supported by start-up funds of Dr. Salvemini and Dr. Arnatt, the National Institutes of Health (NIH) Grant R01NS128004 (to DS and CKA) and the NIH National Institute of General Medical Sciences (Grant T32 GM008306-01) (to KDO). Funding for transcriptomics data was provided by the Intramural Research Program of the National Institutes of Health Clinical Center (ZIACL090034-09, ZIACL090035-08, ZIACL0033-09 to AJM), with supplementary funding provided by the Office of Behavioral and Social Science Research. The contributions of the NIH author(s) were made as part of their official duties as NIH federal employees, are in compliance with agency policy requirements, and are considered Works of the United States Government. However, the findings and conclusions presented in this paper are those of the author(s) and do not necessarily reflect the views of the NIH or the U.S. Department of Health and Human Services.

---

**Supplemental Table 1:** Catalogue and lot numbers for Advanced Cell Diagnostic in situ hybridization probes. The full details of the probes used in multiplex in situ hybridization of human skin samples.

| Gene name | Symbol | Reference no. |
| --- | --- | --- |
| Lysozyme | *LYZ* | 421441 |
| G protein-coupled receptor 183 | *GPR183* | 458801-C2 |
| CD3 epsilon subunit of T-cell receptor complex | *CD3E* | 553971-C3 |
| Interleukin-8 | *IL8* | 310381-C4 |

## Notes

### Competing Interest Statement

The authors have declared no competing interest.

