## Supplementary Figures for "Targeting GPR183 to reduce peripheral sensitization: evidence from rodent and human tissue analyses"

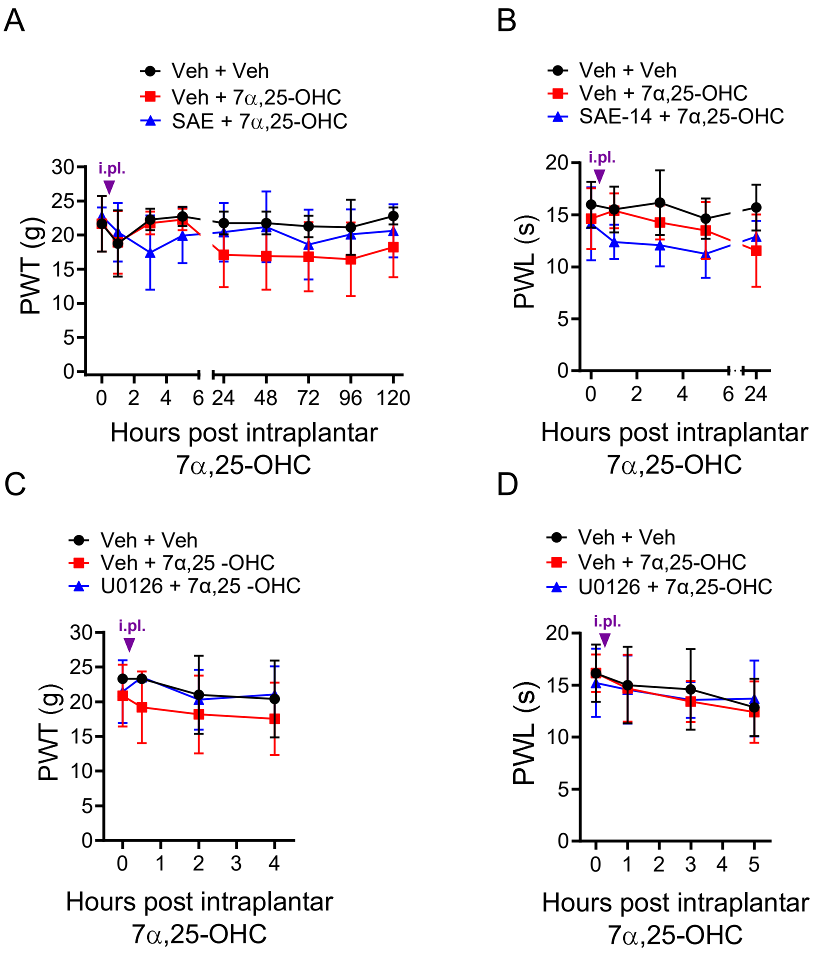


**Supplementary Figure 1: 7α,25-OHC, SAE-14, and U0126 have no effect on mechanical allodynia and thermal hyperalgesia in contralateral paws.**

**A,B)** Intraplantar injections of 7α,25-OHC (5 ng) and SAE-14 have no effect on mechanical allodynia **(A)** and thermal hyperalgesia **(B)** of contralateral paws in male and female rats. **C,D)** 7α,25-OHC-induced (5 ng) and U0126 (10 μg) have no effect on contralateral mechanical allodynia **(C)** and thermal hyperalgesia **(D)**. Data are expressed as mean ± SD for n=6/group **(A,C,D)** or n=4/group **(B)** and analyzed by two-tailed, two-Way ANOVA with Bonferroni multiple comparisons: *$p<0.05$ vs 0 hours; †$p<0.05$ vs. Veh+7*α*,25-OHC group.

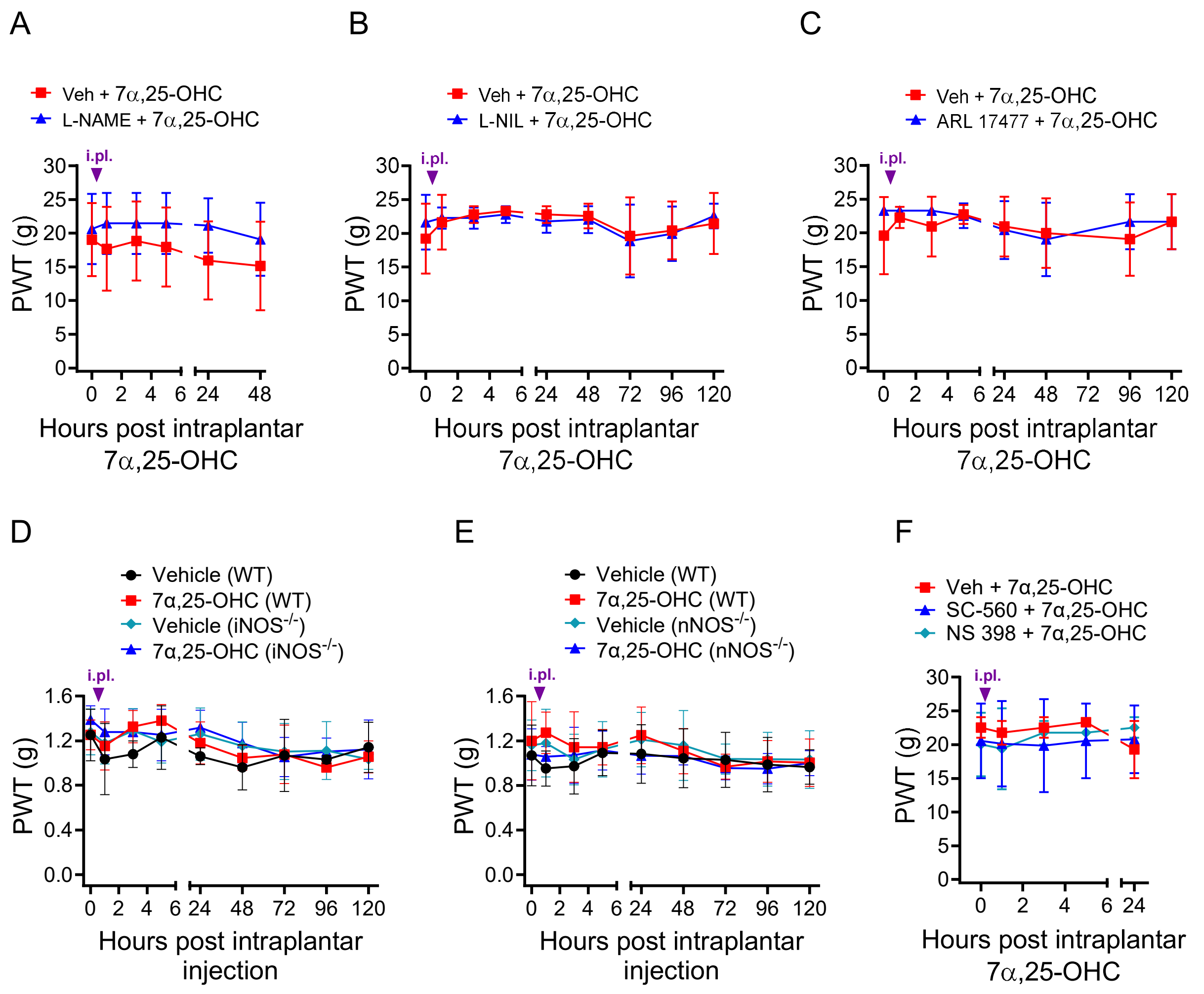


**Supplementary Figure 2: NOS and COX inhibitors have no effect on contralateral paw sensitivity.**

**A, B, C)** Intraplantar injections of 7α,25-OHC (5 ng), L-NAME (50 µg) **(A)**, L-NIL (30 µg) **(B)**, and ARL 17477 (20 µg) **(C)** have no effect on contralateral mechanical sensitivity in male rats. **D, E)** Intraplantar injection of 7α,25-OHC (1 ng) did not elicit contralateral mechanical sensitivities in male WT, iNOS KO **(D)**, and nNOS KO mice **(E)**. **F)** Intraplantar injections of NS-398 (10 µg) or SC-560 (10 µg) had no effect on 7α,25-OHC-induced (5 ng) contralateral mechanical allodynia in rats. Data are expressed as mean ± SD for n=6/group **(A, B, C)**, n=4/group **(D, F)**, or n=7/group **(E)** and analyzed by two-tailed, two-Way ANOVA with Bonferroni multiple comparisons: *$p<0.05$ vs 0 hours; †$p<0.05$ vs. Veh+7*α*,25-OHC group.

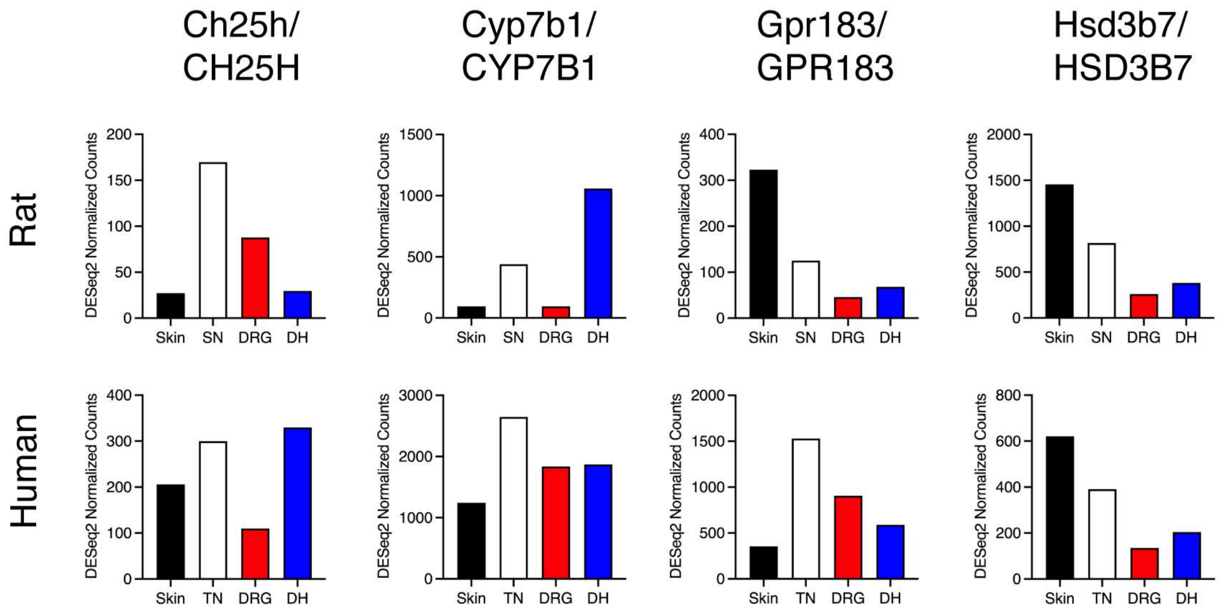


**Supplementary Figure 3. Expression of biosynthesis and degradative enzymes for production, clearance, and signaling of 7-alpha-OHC along the pain neuroaxis.**

A customized RNA-Seq database was uniformly aligned to examine levels of expression of the major enzymes and receptor, GPR183 for 7-alpha-OHC. Biosynthetic enzymes for 7-alpha-OHC were detected along the neuroaxis in the early nociceptive inputs including skin, peripheral nerve (SN in rat, TN in human), sensory ganglion (DRG), and dorsal spinal cord (DH). GPR183 was also present in all four tissues of rat and human. The enzymes responsible for the first step of hydroxylation (biosynthesis) were more prevalent in peripheral nerve as compared to skin in rat, suggesting that the peripheral nerve may be implicated in generation of 7-alpha-OHC, particularly in rat.

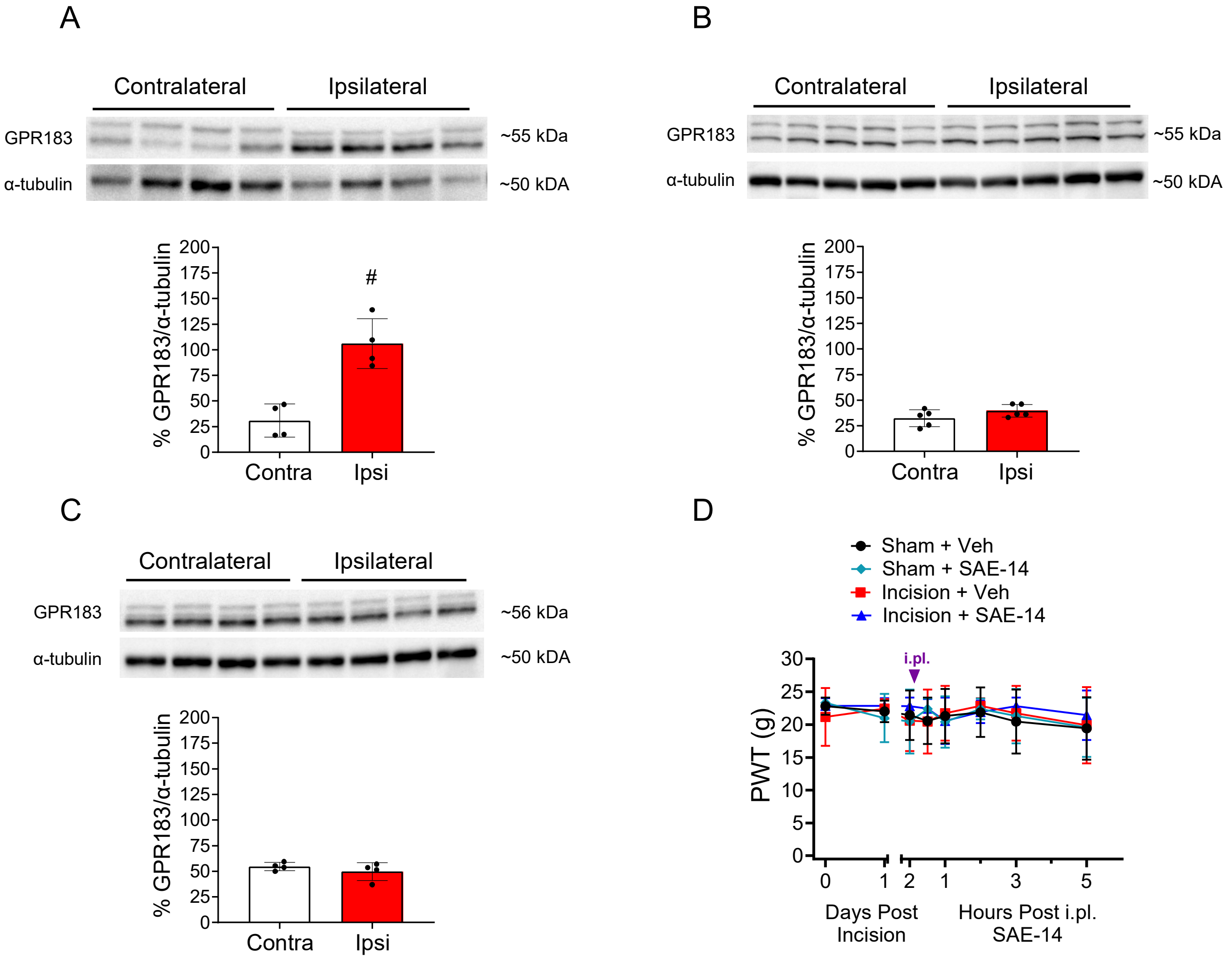


**Supplementary Figure 4. Paw incision injury does not affect GPR183 expression in DRGs or contralateral sensitivity.**

Male and female rats received an incisional injury or sham on one paw and an injection of SAE-14 on Day 2 post op **(Figure 3).** Contralateral paw received no surgery or injections. **A)** GPR183 protein expression was elevated in incised paw tissues from second set compared to naïve paw. **B, C)** L4-L6 dorsal root ganglia in male and female rats day 3 post-hind paw injection show no changes in GPR183 expression in the first set **(B)** or second set **(C)**. **D)** Contralateral paw mechanical allodynia (von Frey) was unchanged following surgery or intraplantar treatment with SAE-14 (600 ng) or its vehicle. Data are expressed as mean ± SD for n=4/group **(A,C)**, n=5/group **(B)**, or n=7/group **(D)** and analyzed by **(A-C)** two-tailed, unpaired t-test, #$p<0.05$ vs Contra, or **(D)** two-tailed, two-way ANOVA with Bonferroni comparisons: *$p<0.05$ vs 0 hours; †$p<0.05$ vs. incision + Veh group.

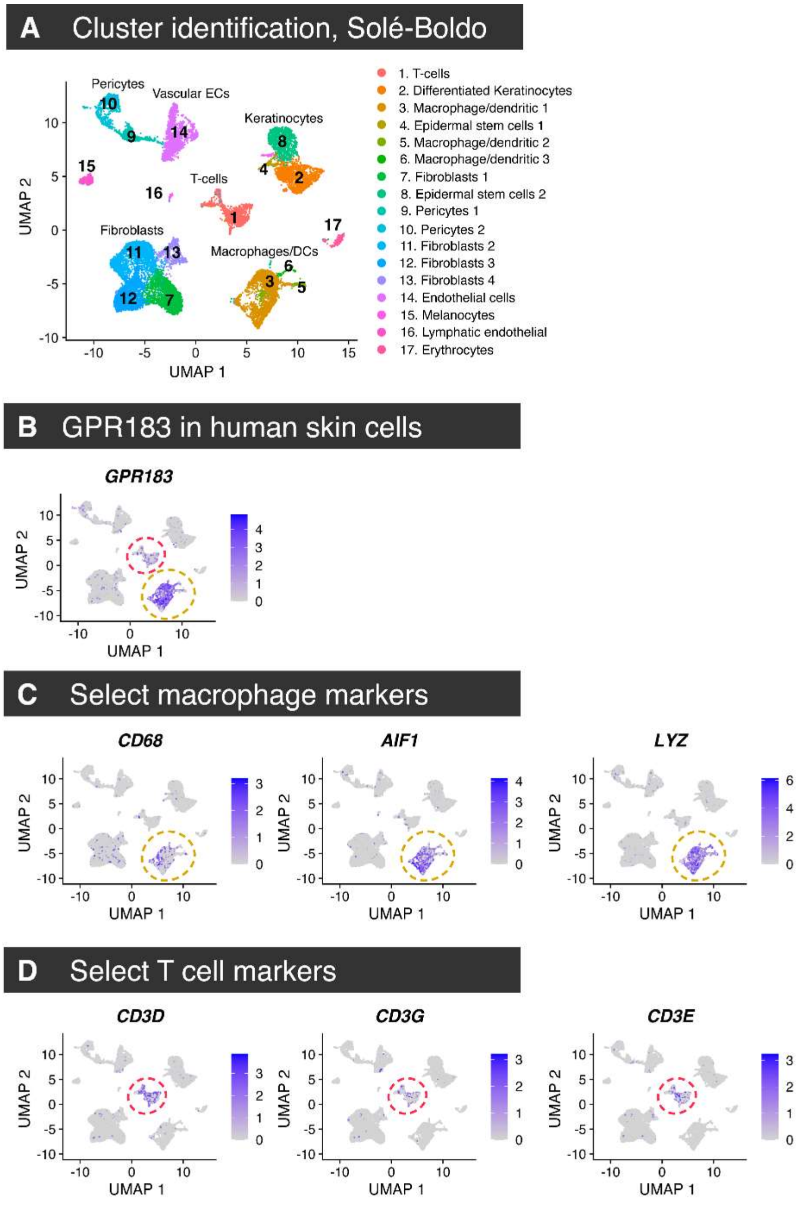


**Supplementary Figure 5. Single cell RNA-Seq analysis of human skin cells expressing *GPR183*.**

To identify the major cell types expressing GPR183 in human skin, a bioinformatic analysis was performed. **A)** Data from a previously published single-cell dataset (PMID: 32327715) was queried and the clusters were adopted from the publication. **B)** For *GPR183* expression was predominantly identified in the clusters corresponding to macrophage/dendritic cells (yellow dashed circle) and T cells (red dashed circle). **C)** As a control, three of the most well-characterized macrophage markers from the literature (*CD68*, *AIF1*, and *LYZ*) are shown, with enrichment in the macrophage cluster, as expected. **D)** Similarly, three of the most highly significant T cell markers (*CD3D*, *CD3G*, and *CD3E*) are shown, with enrichment in the T cell cluster, as expected.

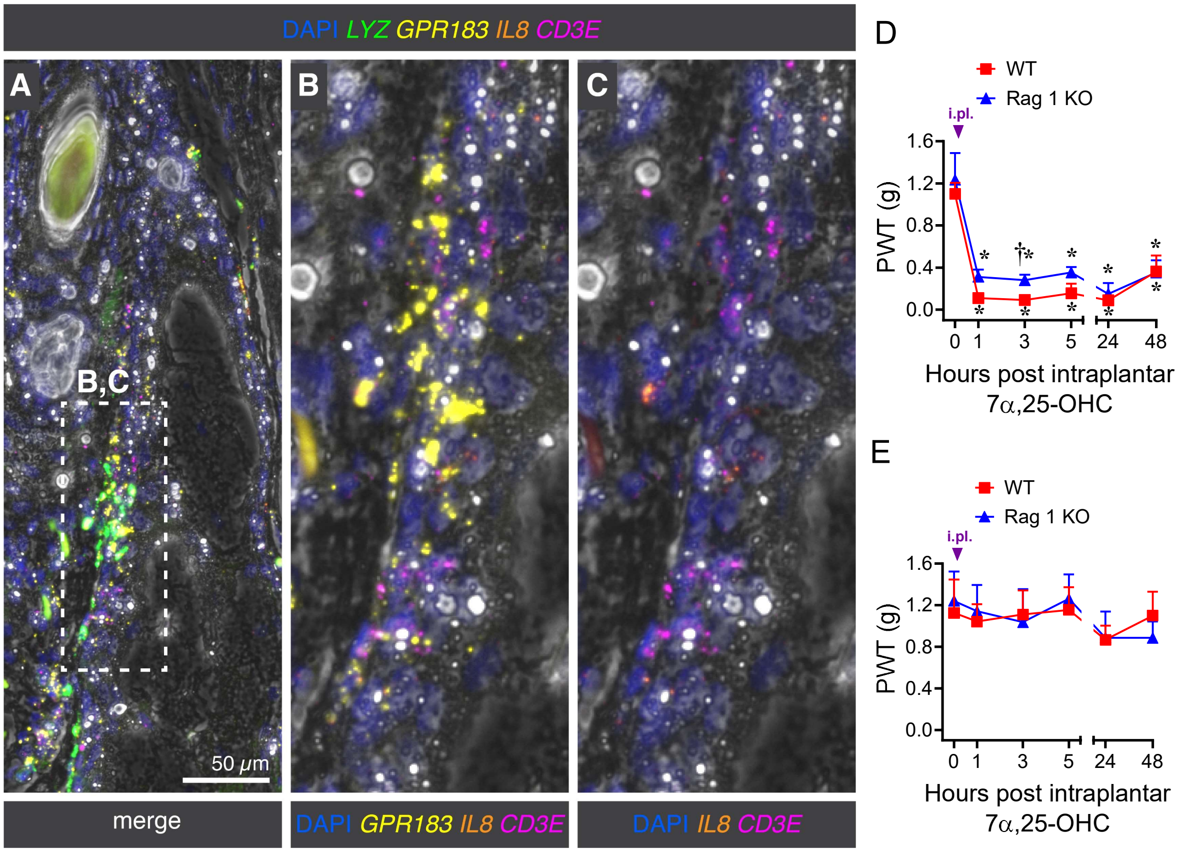


**Supplementary Figure 6. T cells have little-to-no effect on GPR183-mediated hypersensitivity.**

**A)** Enhanced image showing staining for *CD3E* in human skin region adjacent to a perifollicular sebaceous gland. The broad marker of T-cells *CD3E* was weakly detected in human skin, but was identified in an area labeled densely for *GPR183* and *LYZ*$^{+}$ cells adjacent to a perifollicular sebaceous gland. **B, C)** Co-expression was ambiguous as CD3E expression is weak in this structure, but generally did not associate with GPR183 supporting a more prominent role for GPR183 in *CD3E*$^{-}$ populations of cells in human skin. **D, E)** Intraplantar injection of 7α,25-OHC (1 ng) induced mechanical allodynia in WT and Rag 1 KO mice on the ipsilateral side **(D)** with no effect on the contralateral side **(E)**. **D, E)** Data are mean ± SD for n=4/group and analyzed by two-tailed, two-Way ANOVA with Bonferroni comparisons; *$p$<0.05 vs 0 hours; †$p$<0.05 vs. WT.
